# Response to anti-angiogenic therapy is affected by AIMP protein family activity in glioblastoma and lower-grade gliomas

**DOI:** 10.1101/2025.03.13.643116

**Authors:** Humaira Noor, Yuanning Zheng, Haruka Itakura, Olivier Gevaert

## Abstract

**Background:** Glioblastoma (GBM) is a highly vascularized, heterogeneous tumor, yet anti-angiogenic therapies have yielded limited survival benefits. The lack of validated predictive biomarkers for treatment response stratification remains a major challenge. Aminoacyl tRNA synthetase complex-interacting multicomplex proteins (AIMPs) 1/2/3 have been implicated in CNS diseases, but their roles in gliomas remain unexplored. We investigated their association with angiogenesis and their significance as predictive biomarkers for anti-angiogenic treatment response.

**Methods:** In this multi-cohort retrospective study we analyzed glioma samples from TCGA, CGGA, Rembrandt, Gravendeel, BELOB and REGOMA trials, and four single-cell transcriptomic datasets. Multi-omic analyses incorporated transcriptomic, epigenetic, and proteomic data. Kaplan-Meier and Cox proportional hazards models were used to assess the prognostic value of AIMPs in heterogeneous and homogeneous treatment-groups. Using single-cell transcriptomics, we explored spatial and cell-type-specific AIMP2 expression in GBM.

**Results:** AIMP1/2/3 expressions correlated significantly with angiogenesis across TCGA cancers. In gliomas, AIMPs were upregulated in tumor vs. normal tissues, higher- vs. lower-grade gliomas, and recurrent vs. primary tumors (p<0.05). Upon retrospective analysis of two clinical trials assessing different anti-angiogenic drugs, we found that high-AIMP2 subgroups had improved response to therapies in GBM (REGOMA: HR 4.75 [1.96–11.5], p<0.001; BELOB: HR 2.3 [1.17–4.49], p=0.015). AIMP2-cg04317940 methylation emerged as a clinically applicable stratification marker. Single-cell analysis revealed homogeneous AIMP2 expression in tumor tissues, particularly in AC-like cells, suggesting a mechanistic link to tumor angiogenesis.

**Conclusions:** These findings provide novel insights into the role of AIMPs in angiogenesis, offering improved patient stratification and therapeutic outcomes in recurrent GBM.

## Introduction

Angiogenesis is a key pathological hallmark of solid tumors. Among all solid tumors, glioblastomas have the most expansive vascular systems supporting tumor growth and proliferation [1–4]. A range of anti-angiogenic therapies, including anti-VEGF therapies, have been largely explored in treating glioblastoma, and higher-grade astrocytoma/oligodendroglioma during the past decade. Bevacizumab, an anti-VEGF drug was approved by FDA for the treatment of recurrent glioblastoma in 2017 [5]. Although a prolonged progression-free survival was noted, the studies failed to observe any effects on the overall survival of recurrent glioblastoma patients [5–8]. As first-line treatment for newly diagnosed glioblastoma, no survival benefit with the treatment of Bevacizumab was observed [9, 10]. Clinical trial outcomes of other anti-angiogenic therapies such as Aflibercept [11], Dovitinib [12], Sunitinib [13], Sorafenib [14], Cediranib [15], Imatinib [16], Pazopanib [17] and Marizomib [18], were not encouraging for the treatment of recurrent glioblastoma with no observed effects on progression-free survival and/or overall survival. Despite the failures of anti-angiogenic therapies in improving glioma survival, angiogenesis remains an important therapeutic target in this disease due to its potential to treat the highly vascularized tumor. This is substantiated by over forty currently on-going clinical trials of anti-angiogenic drugs for glioblastoma (clinicaltrials.gov).

It is likely that the lack of effective patient stratification and selection has contributed to the limited success of anti-angiogenic therapies in glioblastoma/other higher-grade gliomas, and identifying subgroups that may benefit from these treatments could enhance therapeutic outcomes. Currently, there are no validated biomarkers for effectively selecting glioblastoma patients who would benefit from anti-angiogenic treatments. Thus, predictive molecular biomarkers can be crucial to patient selection, and, ultimately, success in treatment efficacy [19].

Aminoacyl tRNA synthetase complex-interacting multicomplex protein (AIMP) 1/2/3 are parts of a group of multi-functional proteins that form the Aminoacyl-tRNA synthetases (ARSs). The AIMPs are closely linked auxiliary proteins [20] playing a crucial scaffolding role in multi-tRNA synthetase complex assembly [21]. The AIMPs have also shown to play a role in neurological diseases and nervous system functions [22–26], however, despite their involvement in the CNS functions, the genomic and epigenomic roles of the AIMPs have not yet been explored in gliomas. The association between AIMP1 and angiogenesis have been established [27–29], while AIMP2 may play a role in angiogenesis through modulation of the WNT/B-catenin signaling pathway [30], regulating TRAF2 [31], and FUBP1[32] that in turn affect VEGF and angiogenesis. AIMP3 regulates p53 activation and genomic stability [33, 34] that modulates angiogenesis. Despite the relationships between AIMPs and angiogenesis, the genomic and epigenomic role of AIMPs have not yet been studied in light of angiogenesis in cancer. Given the potential theoretical effect of AIMPs in glioma angiogenesis, investigating the influence of AIMPs in glioma angiogenesis and anti-angiogenic treatment response is warranted, particularly in order to explore the potential of AIMPs in stratifying responders to these treatments.

In this study, we comprehensively investigated the association between AIMP expressions and angiogenesis gene-sets across 33 cancers to uncover their link with the angiogenesis pathway. We then focused on differential multi-omic expressions of AIMPs in glioma versus normal brain to gauge the potential impact of AIMP expressions in regulating anti-angiogenic treatment response in glioma. We then analyze the prognostic roles of AIMPs in multi-cohort gliomas regardless of treatment received to provide rationale for retrospectively studying their prognostic roles in two anti-angiogenic treatment specific clinical trial cohorts. We explored AIMPs as predictive biomarkers for anti-angiogenic treatment response in these clinical trials. To identify CpG sites that both regulate AIMPs expression and determine treatment response, that may be used practically at the lab setting to stratify patients, we performed a comprehensive epigenetic study of AIMP CpGs. To provide a possible mechanistic explanation for AIMPs relationship with angiogenesis, we utilized four additional single-cell transcriptomic cohorts, including one spatial transcriptomic cohort, and explored AIMP associations with certain cell-types at single-cell level. In summary, this study aims to lay a foundation for utilizing AIMP status as a potential biomarker for future anti-angiogenic clinical trials.

## Results

### AIMP1/2/3 mRNA expression correlates with angiogenesis pathway across majority of TCGA cancers

To establish possible associations between AIMPs and angiogenesis in cancer, we investigated the correlation between AIMP1/2/3 and the angiogenesis Panther pathway and KEGG pathway gene sets mRNA expressions across 33 TCGA cancers (**Fig.1A-B**). Strikingly, AIMP1 expression showed significant positive correlation across all cancers except CHOL. AIMP2 and AIMP3 expressions showed significant correlation in 24/33 TCGA cancers in Panther pathway analysis (**Fig.1A**). In KEGG pathway gene-set analysis, AIMP2 and AIMP3 expressions showed significant correlation with angiogenesis gene-set expressions in 22/33 and 24/33 TCGA cancers, respectively (**Fig.1B**). These findings strongly suggest an association between AIMP1/2/3 genes and angiogenesis pathways across cancers, providing rationale for investigating the role of AIMPs in anti-angiogenesis treatment response in GBM and LGG due to AIMPs’ role in CNS functions, and the increased vascularity in these tumors.

**Figure 1.**
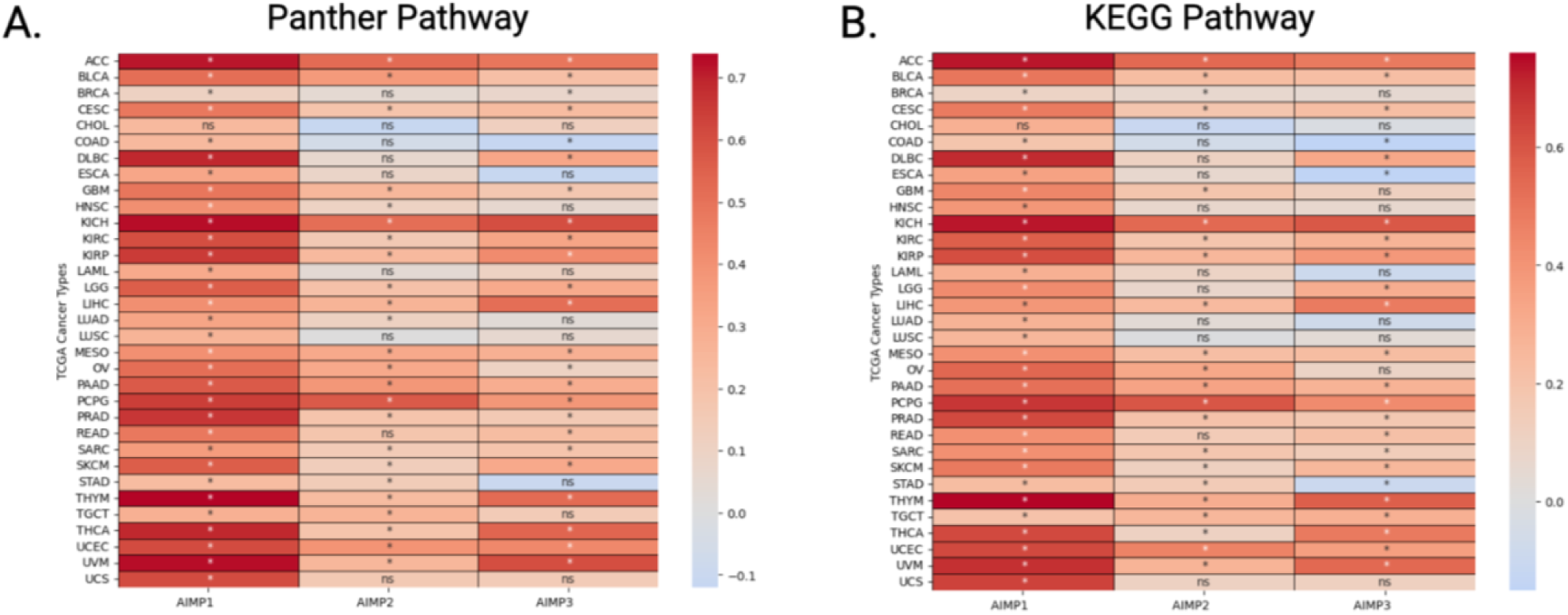
AIMP1/2/3 correlates with angiogenesis pathway gene-sets. Correlation between AIMP1/2/3 mRNA expression and **(A)** 141 Panther pathway angiogenesis gene set and **(B)** 76 KEGG pathway angiogenesis gene set. The color map represents R value. *asterisk represents statistically significant correlation (Pearson’s correlation p<0.05). N.S represents statistically insignificant cases.

Furthermore, to demonstrate that this observed significant association is not normal tissue-specific, but in fact cancer-specific, we compared AIMPs mRNA expression levels in the 33 TCGA cancers with their respective normal tissues (**Supplementary Fig. 1**). The AIMPs were significantly differentially expressed between tumor and normal tissues across various cancers (**Supplementary Fig. 1**). These findings suggest that AIMPs may be involved in common cancer pathways across multiple cancers, including angiogenesis as shown in **Fig.1**, and cancer-type specific investigations of AIMPs are warranted.

### AIMPs mRNA and proteins are differentially expressed in glioblastoma and low grade glioma tissues compared with normal brain tissue

Further, we focused on gliomas and expanded our multi-omic analysis to include protein expression levels to understand the varying correlation results for gliomas in **Fig.1**. First, we investigated the mRNA expression levels of AIMP1/2/3 in TCGA-GBM (n=163), TCGA-LGG (n=512) and normal brain (GTex, n=207) in order to determine possible differential mRNA expression in the tumor tissue compared with normal brain tissue. We found significantly higher mRNA expression levels of AIMP1 and AIMP2 in both GBM and LGG compared with normal brain tissue (**Fig. 2A**, p<0.05 for both). AIMP3 expression was comparable between the tumor tissues and normal brain tissue with no significant difference (p>0.05). At the protein level, however, all three genes showed higher protein expression in immunohistochemical staining (IHC) of GBM tissues and LGG tissues compared with normal brain tissue (**Fig. 2B & 2C**). The staining for AIMP1/2/3 in normal tissue were “low” with weak/moderate intensity, whereas the staining for GBM and LGG were consistently “high” or “medium” with strong/moderate intensity for all three proteins (**Fig. 2C**). This suggests that although at the mRNA level no significant difference in expression was observed for AIMP3 while comparing tumor tissues with normal tissue, AIMP3 may be involved in tumorigenesis and it may be an important factor to investigate in GBM and LGG since an elevated protein expression was observed in IHC. Hence, we included AIMP3 in the analysis of this study.

**Figure 2.**
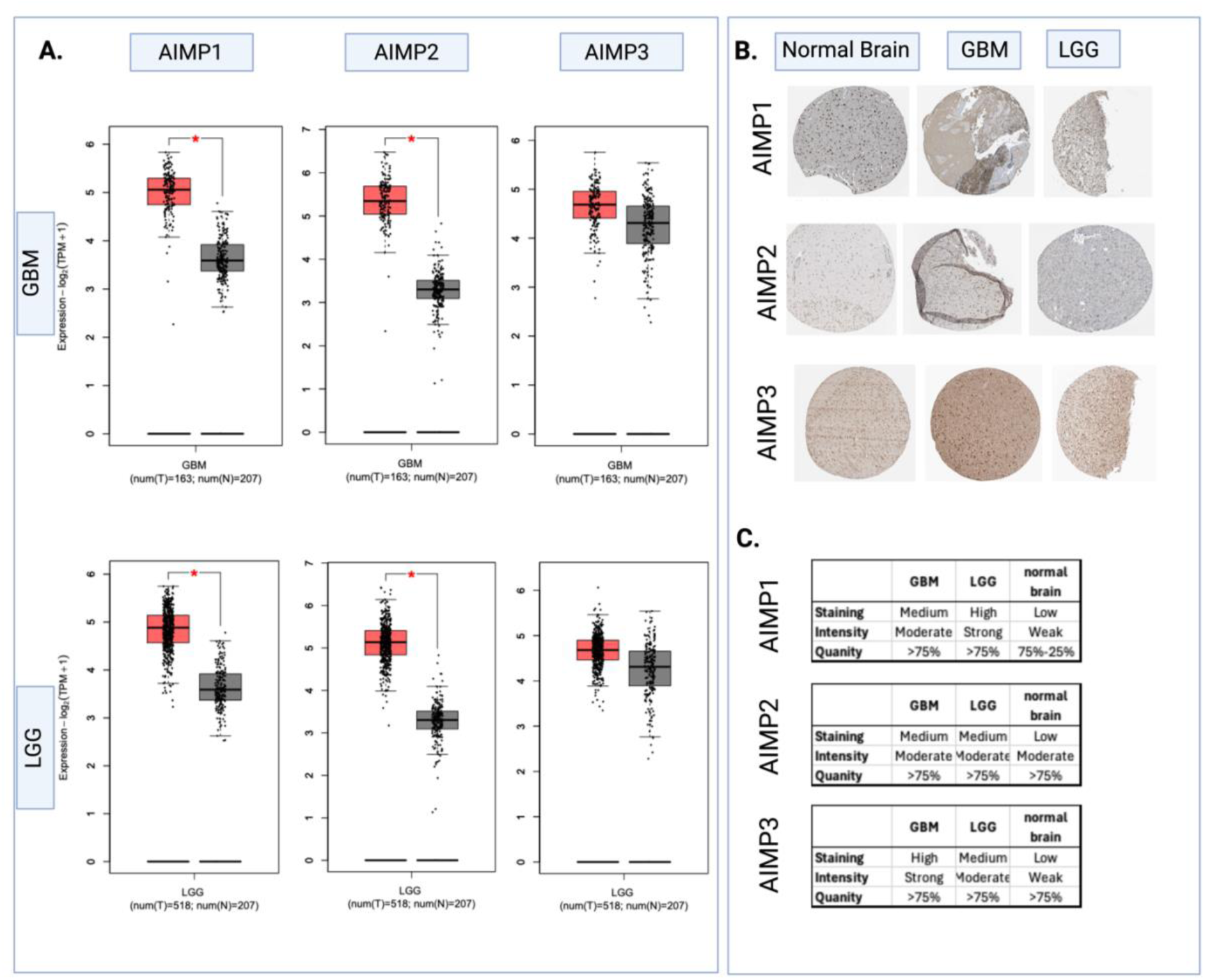
AIMP1/2/3 are differentially expressed in glioma tissues. AIMP1/2/3 (A) mRNA and (B & C) protein expression in TCGA-GBM, TCGA-LGG and normal brain (GTex) tissues. Data for (B) and (C) were collected from the Human Protein Atlas. Student’s t-test * p<0.05.

### AIMPs are associated with higher tumor grade and recurrence, and their prognostic effects may potentially be treatment-dependent

To understand the association between AIMPs and tumor aggressiveness, we compared AIMPs mRNA expression levels between low- (grade II) and high- (grade II & IV) tumors (**Fig.3Ai**). AIMP1 and AIMP2 were significantly upregulated in high-grade tumors compared with low-grade tumors. We compared AIMPs expression between primary and recurrent tumors to understand whether AIMPs are recurrence-related and may be of particular importance in tumor progression (**Fig3. A-ii**). We found that all three AIMPs were significantly upregulated in recurrent gliomas compared with primary gliomas. This observation supports the association between tumor recurrence/progression and increased AIMP levels, and substantiates the basis for investigating AIMPs particularly in recurrent GBM.

Before investigating AIMPs’ prognostic role in anti-angiogenic therapy treated patients, we analyzed their prognostic role in multiple glioma cohorts with heterogeneous treatment. This can potentially indicate any role AIMP1/2/3 may play in tumor aggressiveness and progression in general, regardless of treatment. We performed survival analysis based on AIMP1/2/3 mRNA expression levels in four GBM cohorts (TCGA n= 151, CGGA n=220, REMBRANDT n=181, and Gravendeel n=155), and five LGG cohorts (TCGA-astrocytoma n=193, TCGA-oligodendroglioma n=99, CGGA-astrocytoma n=88, REMBRANDT-oligodendroglioma n=50, and REMBRANDT-astrocytoma n=104). The Kaplan-Meier survival curves stratified by high- and low-AIMP1/2/3 mRNA levels are presented in **Figure 3B** for GBM and **Supplementary Figure 2** for LGG.

**Figure 3.**
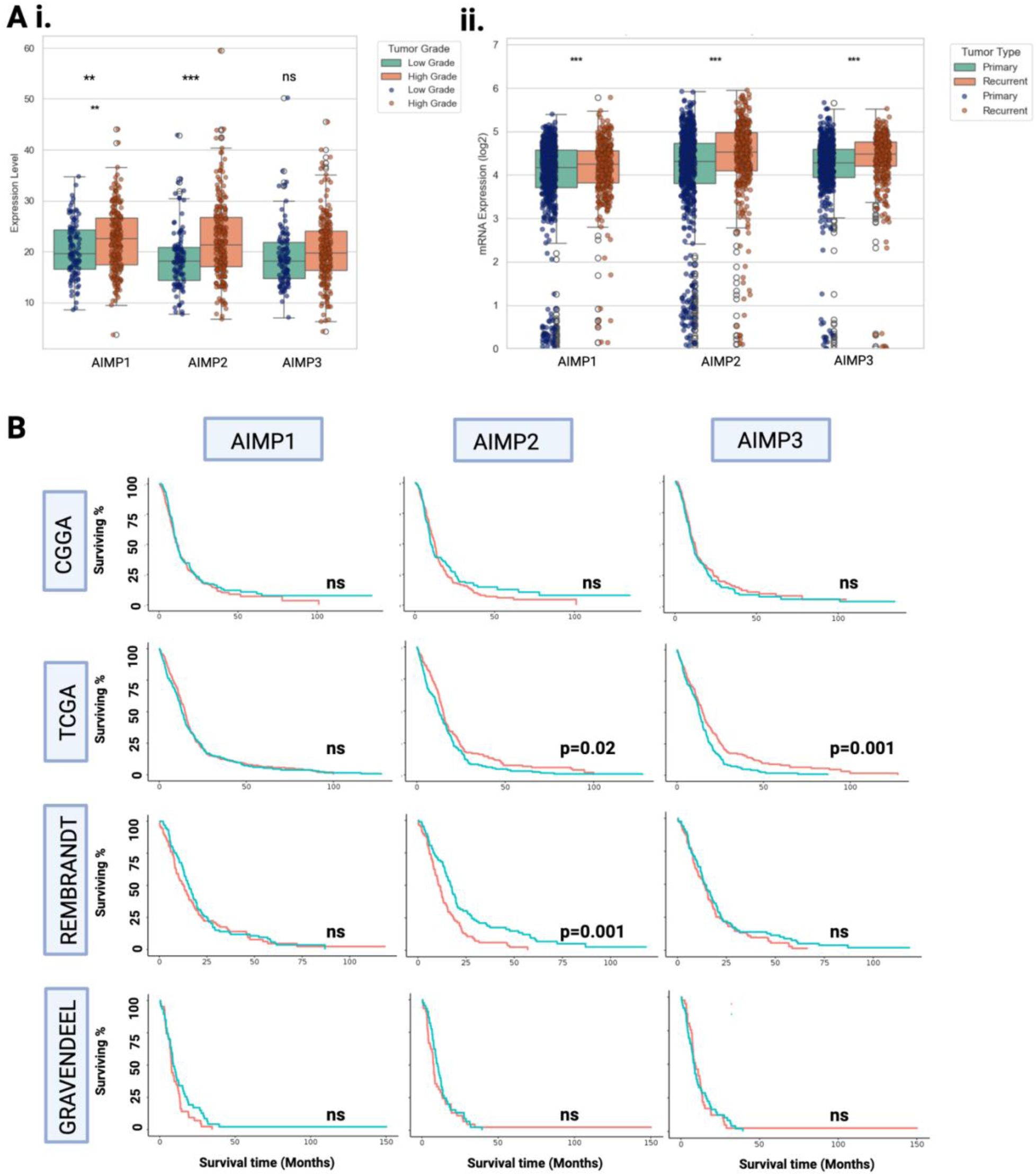
AIMPs are associated with tumor aggressiveness, recurrence and prognosis. AIMPs mRNA expression levels in **(Ai)** low (II; n=103) and high-grade (III & IV; n=218) gliomas, and **(Aii)** primary (n=229) and recurrent gliomas (n=62) in CGGA dataset. Student’s t-test p-value<0.05 considered statistically significant. **(B)** Kaplan-Meier Survival Curves depicting prognostic effects of AIMP1/2/3 mRNA expressions in CGGA, TCGA, REMBRANDT and Gravendeel cohorts of GBM. Red=high-expression green=low-expression based on median-cutoff; log-rank p-value<0.05 is considered significant.

In GBM, AIMP1 was not found to be a significant prognostic factor in any of the four cohorts. There were inconsistent results for AIMP2 in TCGA and REMBRANDT cohorts, where AIMP2 expressions significantly affected prognosis, however, in TCGA high expression was associated with improved survival while in REMBRANDT it was associated with poorer survival (p<0.05; **Fig.3B**). AIMP3 showed significant prognostic relevance in only the TCGA cohort, with high expression levels associating with improved survival (p<0.05; **Fig.3B**). The results for LGG were of similar patterns (**Supplementary Fig.2**), with no corroborating results across cohorts.

Our analysis suggests that the prognostic roles of AIMPs expression are cohort-dependent, and are not consistent across multiple cohorts. A likely explanation for this is the possible heterogenous treatment received by patients in each cohort, and the possibility that AIMPs expression potentially affects treatment response. This observation is potentially aligned with the hypothesis of our study and supports further investigations into the role of AIMPs in treatment response in homogeneously treated cohorts, particularly anti-angiogenic therapies.

### Anti-angiogenic treatment shows efficacy in highly expressed AIMP subgroups of GBM in homogeneously treated REGOMA and BELOB clinical trials

To understand whether AIMPs have a role in determining anti-angiogenic treatment response, we retrospectively analyzed the outcome of two GBM clinical trials with homogeneous treatment, REGOMA [35, 36] and BELOB [37], where pre-treatment transcriptomic data were publicly available. We investigated overall survival in patients following anti-angiogenic treatments within AIMP1/2/3 high- and low-expression sub-groups (**Fig. 4**). The REGOMA trial, an Italian clinical trial for recurrent GBM patients following chemoradiation therapy, compares the efficacy of CCNU (i.e Lomustine) vs Regorafenib (an anti-angiogenic drug) monotherapy. In the original study Regorafenib was associated with significantly improved OS compared with CCNU monotherapy, however, our retrospective analysis of this cohort on the basis of AIMP1/2/3 expression subgroups revealed that this association is only observable in AIMP1/2/3 high expression subgroups (p<0.05; **Fig. 4**), and not in the low expression subgroups (p>0.05; **Supplementary Figure 3**). Regorafenib treatment was associated with significantly improved survival compared with CCNU treatment in high AIMP1 (HR[95% CI]: 2.41[1.15-5.06]; p=0.019), high AIMP2 (HR[95% CI]: 4.75[1.96-11.5]; p<0.001), and high AIMP3 (HR[95% CI]: 2.45[1.14-5.27]; p=0.021) expression subgroups (**Fig.5A**).

**Figure 4.**
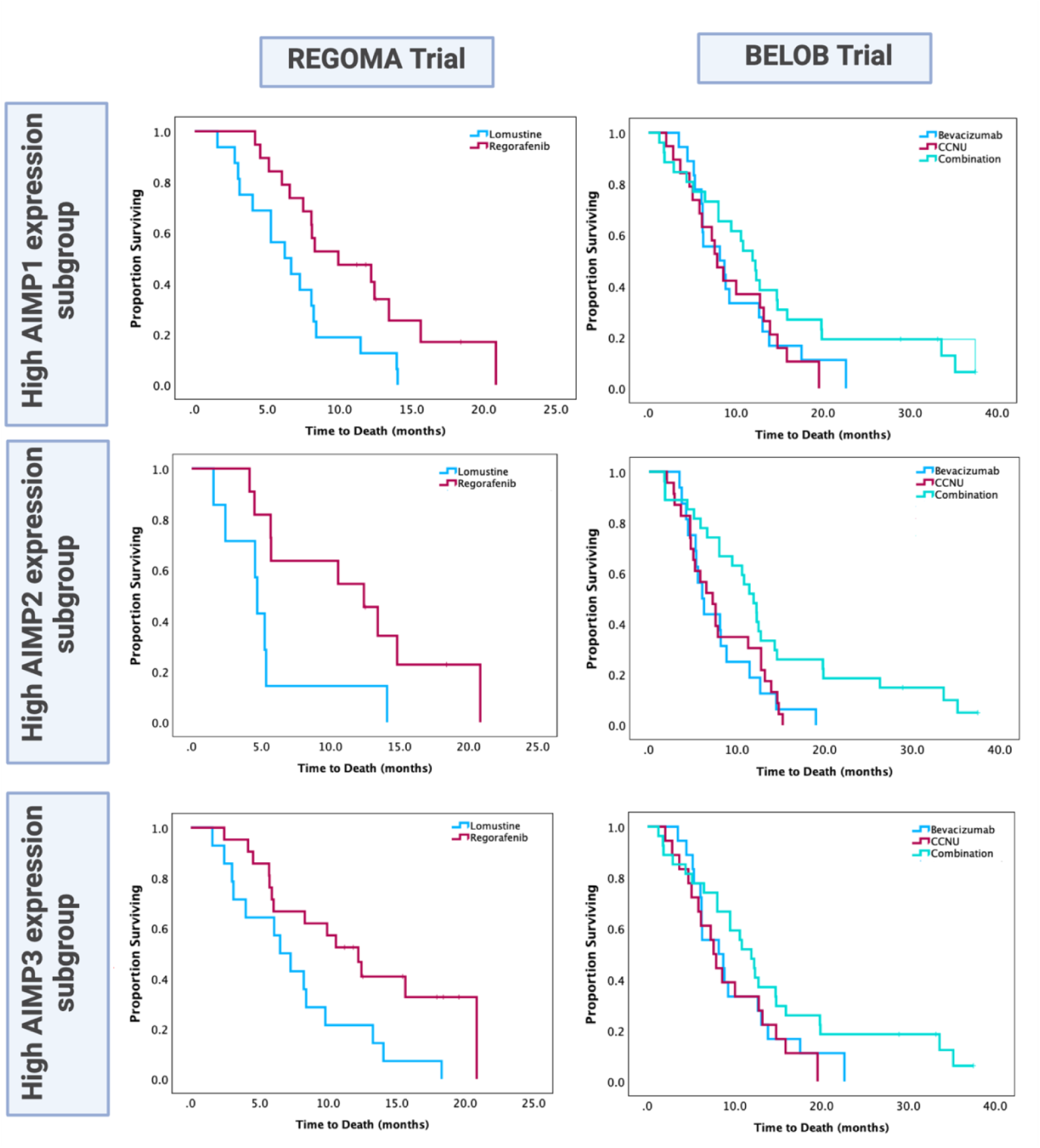
High AIMP mRNA expression subgroups are more responsive to anti-angiogenic therapies. Kaplan-Meier survival analysis on retrospective clinical trials of recurrent GBM (REGOMA and BELOB trials). High expression sub-groups are stratified by median mRNA expressions of AIMP1/2/3. Log-rank p-value<0.05 is considered significant.

**Figure 5.**
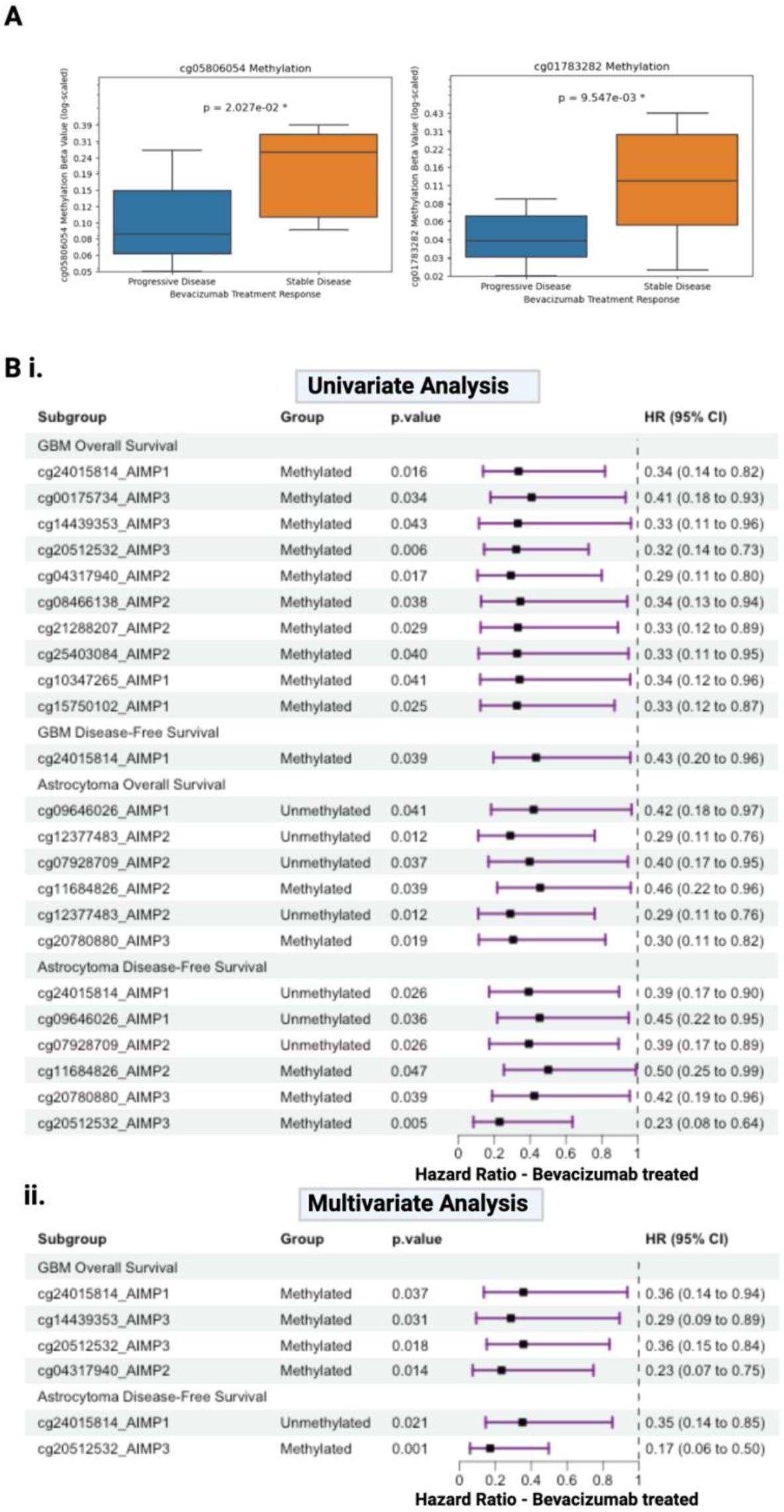
Bevacizumab treatment response is associated with specific AIMP CpG site methylation status in gliomas. **(A)** Significant association between two CpG sites and astrocytoma (n=22) pathological response to bevacizumab treatment. Student’s t-test p-value<0.05 considered significant **(B)** Forest plots depicting Cox proportional hazards model univariate **(i)** and multivariate **(ii)** analysis for the effects of AIMP1/2/3 CpG methylation status

The BELOB trial was a Netherlands-based clinical trial for 148 recurrent GBM patients following chemoradiation. The BELOB trial compares treatment with bevacizumab, CCNU (i.e Lomustine), and combination therapy with the two [37]. In the original BELOB study, no benefit for combination therapy or bevacizumab alone was observed, however, in our retrospective sub-group analysis, we identified significantly improved survival in patients with combination therapy compared with bevacizumab alone (HR[95% CI]: 2.3[1.17-4.49]; p=0.015) or CCNU alone (HR[95% CI]: 1.99[1.084-3.66]; p=0.026) in the AIMP2 high expression sub-group (**Fig. 4**). This was not observed in AIMP1 or AIMP3 high expression sub-groups (**Fig. 4**), or any of the AIMP1/2/3 low expression groups (**Supplementary Figure 3**). This observation underscores the importance of AIMP2 as a predictive biomarker for anti-angiogenesis combination treatment response, which was not observed in the mixed group in the original study.

The results from these two retrospective analysis based on AIMP expression sub-groups indicate that AIMPs, particularly AIMP2, may influence response to anti-angiogenic therapies such as bevacizumab and regorafenib in recurrent GBMs on which chemoradiation therapy was ineffective. The type of anti-angiogenic drug was a factor in determining whether anti-angiogenic monotherapy was more efficacious or there was greater benefit with CCNU combination treatment. In both cases, the denominating factor of anti-angiogenic treatment addition improved patient survival in high AIMP2 expression subgroups of recurrent GBM patients. These findings underscore the need for considering AIMP expression status in evaluating response to anti-angiogenic therapy in recurrent GBM, and they support the potential of AIMP status as a biomarker for patient selection in upcoming clinical trials.

### Potential applicability of specific AIMP CpG methylation status as predictive biomarker for anti-angiogenic therapy response

Since AIMP high versus low groups are relative, and no threshold exists for determining potential high-expression subgroups except a relative median-cut off, we sought to identify specific AIMP CpGs whose methylation status may act as a practically applicable threshold for these sub-groups. This is similar to MGMT CpGs (cg12981137, cg12434587, cg12981837, and cg03071809) that determine MGMT methylation status, acting as potential predictive biomarker for chemotherapy response [38]. For this, we firstly correlated AIMPs mRNA expression with AIMPs CpG methylation levels in TCGA cohorts, to identify CpGs that may potentially epigenetically regulate the AIMPs (**Supplementary Figure 4**). The significance of identifying epigenetic regulators of gene expression is well established, as evidenced by the development of numerous bioinformatics tools designed for this purpose [39–41]. After identifying CpGs that significantly correlate with their respective mRNA expression levels, we performed a CpG-wide survival analysis to determine prognostic CpGs regardless of treatment type (**Supplementary Table 1**). AIMP1 CpGs cg20061689 and cg07367189, that do not correlate with mRNA expression, showed significant prognostic effects in GBM OS. AIMP2 CpG cg11684826, that negatively correlate with AIMP2 expression, was a significant prognosticator for both OS and DFS in GBM. This suggests that cg11684826 prognostic effects may not be anti-angiogenic treatment specific, and it may not be a useful predictive biomarker although it showed significant correlation with expression. In LGG, there were no prognosticators for astrocytoma OS, however, AIMP3-cg05806054 and AIMP3-cg01783282 were significantly associated with astrocytoma DFS (**Supplementary Table 1**). These two CpG site methylation levels were also significantly lower in stable disease vs progressive disease in bevacizumab-treated astrocytoma (**Fig.5A**), and may play a role in astrocytoma treatment response. However, they do not correlate with AIMP3 mRNA expression levels in LGG, and therefore the methylation status of these CpGs cannot be used to determine AIMP3 expression sub-groups (**Supplementary Figure 4**).

To identify anti-angiogenic treatment-specific AIMP CpG prognosticators, we examined the OS and DFS of TCGA-GBM and TCGA-LGG where treatment data were available for bevacizumab status. In TCGA-LGG, we only analyzed the astrocytoma subtype since there was limited bevacizumab treatment data for oligodendroglioma. We performed Cox proportional hazards model analysis at the univariate and multivariate levels (adjusting for age, sex, race and grade) and significant results are depicted in the forest plots in **Figure 5Bi-ii**. We identified four AIMP1/2/3 CpG sites in GBM that remained significant prognosticators of OS after adjusting for age, sex, and race (**AIMP1-cg24015814, AIMP2-cg04317940, AIMP3-cg14439353 and AIMP3-cg20512532**). Methylation levels of these CpGs significantly correlate with their respective mRNA expression levels, and may thus be useful in stratifying AIMP high expression subgroups that may benefit from anti-angiogenic therapies in GBM. **AIMP1-cg24015814** and **AIMP3-cg20512532** were also associated with improved DFS with bevacizumab treatment in astrocytoma at multivariate analysis **(Fig. 5**), and may thus be of particular importance in gliomas in general, warranting further investigations which is beyond the scope of this study.

From this analysis, we report four CpG site methylations predictive of bevacizumab response that may be practically utilized at the lab setting to stratify possible responders (**AIMP1-cg24015814, AIMP2-cg04317940, AIMP3-cg14439353 and AIMP3-cg20512532)**. Our collective results from the retrospective clinical trial analysis suggest that AIMP2 expression may be the strongest candidate for predicting anti-angiogenesis response, thus we emphasize on the applicability of **AIMP2-cg04317940** as a stratifying factor. in bevacizumab treatment response in TCGA-GBM (n=80) and TCGA-LGG (n=47). Only significant results reported (Chi-square p-value<0.05.)

### AIMP2 is upregulated in EGFR-Related AC-Like GBM Cells at Single-Cell Resolution with Homogeneous Spatial Distribution

Finally, to investigate the spatial distribution of AIMP2 expression within glioblastoma (GBM) tumors, we analyzed a spatial transcriptomic dataset [42]. This analysis revealed a homogeneous expression pattern of AIMP2 across tumor tissues, with expression levels ranging from low to high but lacking distinct regions of co-localization **(Fig. 6ai–ii)**. The widespread expression of AIMP2 may have significant therapeutic implications for anti-angiogenic treatments, as it indicates that the entire tumor—not just isolated regions—may rely on angiogenesis-driven survival mechanisms. Since our results revealed that AIMP2-high tumors respond favorably to anti-angiogenic therapy, this homogeneous expression pattern may ensure that anti-angiogenic therapy may exert a more uniform therapeutic effect, potentially reducing the likelihood of therapy-resistant subpopulations emerging within the tumor. The expression patterns of AIMP1 and AIMP3 were similar to that of AIMP2 (**Supplementary Fig.5**). It may be useful to note here that with recently developed advanced computational models, AIMP mRNA expressions may be predicted directly from histology slides for patient stratification without the need for RNA sequencing [43].

**Figure 6.**
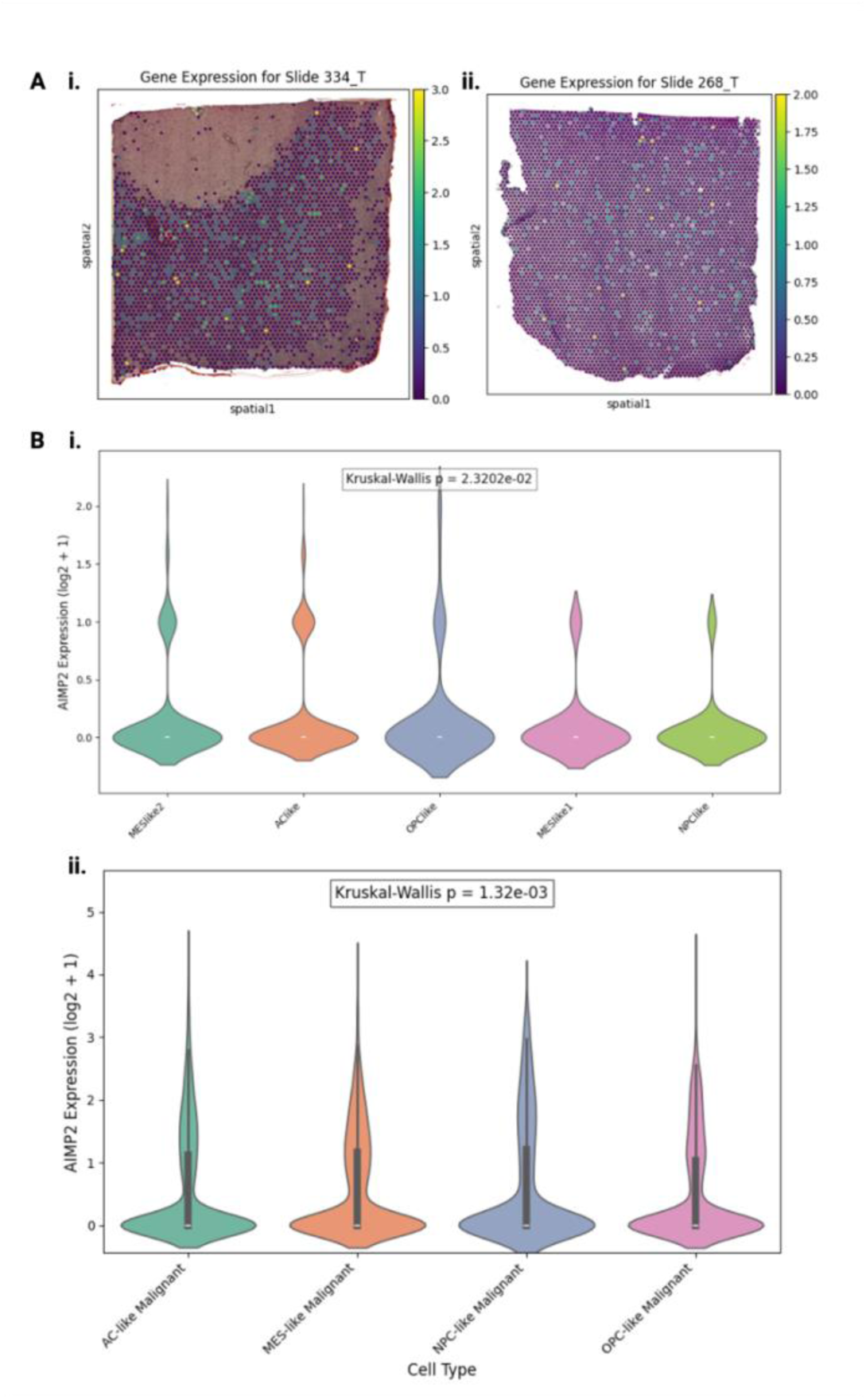
Spatial and cell-type-specific expression of AIMP2 in glioblastoma (GBM) tumors. **(A)** Spatial distribution of AIMP2 expression across tumor tissues in two representative GBM slides (i: Slide 334_T, ii: Slide 268_T) analyzed using a spatial transcriptomic dataset. The heatmaps display homogeneous expression of AIMP2 with varying intensity from low (purple) to high (yellow), indicating the absence of localized expression hotspots. **(B)** Violin plots representing AIMP2 expression across GBM cellular subtypes. **(i)** AIMP2 expression in key GBM subtypes, including MES-like, AC-like, OPC-like, and NPC-like cells from a concatenated single-cell RNA-seq cohort (Kruskal-Wallis p = 2.32e-02). **(ii)** AIMP2 expression across malignant GBM subtypes from a spatial transcriptomic dataset. (Kruskal-Wallis p = 1.32e-03).

To determine the cell-type specificity of AIMP2 expression, we compared its expression levels across key GBM cellular subtypes—MES-like, AC-like, OPC-like, and NPC-like—in both the spatial transcriptomic dataset and a separate concatenated single-cell RNA-seq cohort comprising three independent GBM datasets **(Fig. 6bi–ii)**. In both datasets, AC-like cells exhibited the highest AIMP2 expression, followed by MES-like cells (Kruskal-Wallis p<0.05). Given that AC-like cells are characterized by EGFR amplification/overexpression [44, 45], which regulates VEGF expression [46], the enrichment of AIMP2 in this subtype may explain a possible mechanistic link between AIMP2 expression and the tumor’s angiogenic profile. This connection further supports the observed therapeutic benefit of anti-angiogenic treatment in AIMP2-high tumors.

## Discussion

Anti-angiogenesis therapies have been studied in gliomas for a long time, however, its efficacy has not been reflected through improving the overall survival of the patients. These therapies continue to be tested in new clinical trials to this date as a potential treatment due to the high vascularization in gliomas, particularly GBM. Currently, there is no validated predictive biomarker for stratifying the highly heterogeneous tumors for potential response to anti-angiogenesis. Given the link between AIMP1 and angiogenesis [27–29], and the potential angiogenesis regulation by AIMP2 and AIMP3 through modulation of the WNT/B-catenin signaling pathway [30], and p53 activation [33, 34], respectively, we explored our hypothesis of AIMPs playing a key role in anti-angiogenesis treatment response. In this study, we show that AIMPs expressions’ are associated with angiogenesis pathway genes. This link between AIMPs and angiogenesis pathway has not been previously reported, however, there have been reports of their association with immune-related pathways. Qiu et. al reported in their pan-cancer study that higher AIMP2 expression may be a potential biomarker for breast cancer in the context of immunotherapy response [47]. They focused on the association between immune regulation and tumor immune microenvironment, and did not delve into exploring angiogenesis pathway association. Similarly, AIMP1 was previously reported as part of an 8-gene immune-related signature for GBM [48]. Since VEGF, a key player in angiogenesis, may modulate immune response in glioma [49], these studies support our findings of the angiogenesis pathway association with AIMPs.

Here, we report aberrant protein expressions of AIMP1, AIMP2 and AIMP3 in gliomas compared to normal brain, and their significantly higher mRNA expression in higher grade compared to lower grade tumors. We also established that recurrent GBM tumors have significantly higher expression of AIMP1/2/3 compared to primary GBM tumors. These findings indicate a link between AIMP1/2/3 and tumor aggressiveness and tumor recurrence. Thus, although the focus of this study is anti-angiogenesis therapy, it should be noted that AIMP1/2/3 inhibition may be a potential therapeutic approach that may benefit gliomas. Pre-clinical brain-permeable AIMP2 inhibitors have shown some promise in Parkinson’s disease [50], that may be tested in glioma cell lines in future studies. Currently, there are no reported AIMP1 and AIMP3 inhibitors.

Through the survival analysis of four cohorts, we showed that the prognostic impact of AIMP1/2/3 is potentially treatment dependent, and through retrospective analysis of two recurrent GBM anti-angiogenic clinical trials we established the role of AIMP2 as a predictive molecular biomarker for anti-angiogenic treatment response. We showed that high-AIMP2 mRNA subgroups responded better to two different anti-angiogenic therapies, reflected by improved overall survival. Previously, in a single-institutional study, TMEM173 and FADD were reported as potential biomarkers at proteomic level analysis [51], however, the current study is the first report of an anti-angiogenic treatment biomarker at overall survival level, validated by two recurrent GBM clinical trials.

We report the specific CpG site methylation of AIMP2 cg04317940 as a practically applicable stratifying factor for high-AIMP expression patient subgroups who may benefit from anti-angiogenesis therapy, much like specific CpG site stratifiers of MGMT methylation for predicting temozolomide response at the lab setting [38]. However, it should be noted as a limitation that this CpG was identified based on bevacizumab treatment response in particular, and it should be validated upon availability of methylation data for other anti-angiogenic treatment datasets.

Our single-cell transcriptomic analysis provides a possible explanation for the enhanced efficacy of anti-angiogenic therapy observed in AIMP2-high tumors. AIMP2 was homogenously distributed across GBM tumor tissues without forming localized clusters or regions of concentrated expression. This implies that the tumor may rely on angiogenesis-driven pathways for survival, potentially explaining the observed enhanced response of AIMP2-high tumors to anti-angiogenic therapies with a more uniform therapeutic effect across the tumor. Moreover, we determined that AC-like cells, that are associated with EGFR amplification/overexpression [44, 45], display the highest AIMP2 expression levels across cell-types in two single-cell transcriptomic cohorts. It is known that EGFR regulates VEGF, which is a key player in angiogenesis [46], thus this may be the mechanistic explanation for the role of AIMP2 in modulating angiogenesis treatment response.

Based on our findings, we propose that utilizing AIMP2 high expression as a predictive biomarker, stratified by methylation, for patient selection can potentially enhance therapeutic efficacy and improve outcomes in recurrent glioblastoma patients receiving anti-angiogenic treatments.

## Methods

### Glioma datasets

Multiple publicly available glioma datasets were used for analysis across several sections of this study. These included multi-institutional datasets with genomic and clinical data (TCGA, CGGA, REMBRANDT, and GRAVENDEEL), two anti-angiogenic therapy clinical trials of recurrent GBM (REGOMA and BELOB trials), one spatial transcriptomic GBM dataset [42], and three concatenated single-cell transcriptomic GBM dataset (GSE131928, GSE163108, and GSE84465). In this section, we detail the sources and data modalities for all dataset for reference.

TCGA-GBM (n=152) and TCGA-LGG (n=292) transcriptomic, CpG-level epigenetic and clinical data were downloaded from www.cbioportal.org. For transcriptomic data, only cases with available RNA Sequencing data were selected. While pathological treatment response data for TCGA-GBM was not available, it was available for certain TCGA-LGG (particularly for astrocytoma subtype). Thus, bevacizumab pathological treatment response analysis was performed on only TCGA-astrocytoma.

CGGA-GBMLGG (total n=325) transcriptomic and clinical data were downloaded from www.cgga.org.cn. For transcriptomic data, only cases with available RNA Sequencing data were selected. Primary (n=229) and recurrent gliomas (n=62) were present in this dataset, unlike TCGA where only primary tumors were predominant. REMBRANDT-GBM (n=219), REMBRANDT-LGG (n=225), GRAVENDEEL-GBM (n=159) and GRAVENDEEL-LGG (n=117) transcriptomic and clinical data were downloaded from https://gliovis.bioinfo.cnio.es/. In these cohorts, transcriptomic data available were microarray. All AIMP1/2/3 microarray data were available.

The REGOMA clinical trial [35, 52] (n=71) transcriptomic and clinical data were downloaded from https://www.ncbi.nlm.nih.gov/geo/query/acc.cgi?acc=GSE154041. For this dataset, the transcriptomic data available were processed with RNA sequencing. The BELOB clinical trial [37] (n=112) transcriptomic and clinical data were downloaded from https://www.ncbi.nlm.nih.gov/geo/query/acc.cgi?acc=GSE72951. For this dataset, the transcriptomic data available were processed with microarray. Both of these clinical trials assessed anti-angiogenic treatment for recurrent GBMs.

Single-cell transcriptomic cohorts analyzed included GSE131928, GSE163108, and GSE84465 (Sources: https://www.ncbi.nlm.nih.gov/geo/query/acc.cgi?acc=GSE131928, https://www.ncbi.nlm.nih.gov/geo/query/acc.cgi?acc=GSE163108, and https://www.ncbi.nlm.nih.gov/geo/query/acc.cgi?acc=GSE84465, respectively.)

One spatial transcriptomic cohort [42] (n=16) was used for spatial visualization of AIMP expressions (https://datadryad.org/dataset/doi:10.5061/dryad.h70rxwdmj#readme).

### Pan-cancer association between angiogenesis genes and AIMP1/2/3

Pan-cancer analysis to determine the association between angiogenesis pathway genes and AIMP1/2/3 mRNA expression levels were performed on the 33 TCGA cancers. Normalized pan-cancer transcriptomic data were downloaded from GEPIA2 platform (http://gepia2.cancer-pku.cn/). Two angiogenesis-related gene sets were used for correlation with AIMP expressions across TCGA cancers. First gene set was Panther pathway angiogenesis (141 genes) and the gene set was downloaded from https://maayanlab.cloud/Harmonizome/gene_set/Angiogenesis/PANTHER+Pathways. The second gene set was KEGG pathway angiogenesis (76 genes) and the gene set was downloaded from https://www.gsea-msigdb.org/gsea/msigdb/cards/kegg_vegf_signaling_pathway. For each cancer type, the AIMP1/2/3 mRNA expression levels were correlated with the combined gene-set signature mRNA expression levels. Pearson’s correlation coefficients were reported, with p-value<0.05 considered statistically significant. Heatmaps were created with the Seaborn package using Python.

### Multi-omic differential expression of AIMP1/2/3 in gliomas

Firstly, transcriptomic data were analyzed to understand differential expressions of AIMP1/2/3 in gliomas. The mRNA expression levels (RNASeq) of AIMP1/2/3 in TCGA-GBM and TCGA-LGG were compared with normal brain tissue transcriptomic data from GTEx (normal tissue RNASeq data downloaded from https://gtexportal.org/). Student’s t-test comparison was performed to compare tumor vs normal groups, with p-value<0.05 considered statistically significant.

To compare primary versus recurrent tissues, the mRNA expression levels were analyzed in CGGA data set, where sufficient primary and recurrent glioma cases were available for analysis (refer to “Glioma datasets” section). To compare expression differences between grades, the AIMP1/2/3 mRNA expression levels in lower (WHO grade II) and higher (WHO grade III and IV) grade gliomas were compared in the CGGA glioma dataset. Student’s t-test comparison was performed, with p-value<0.05 considered statistically significant.

To visualize and compare AIMPs at the proteomic level, we analyzed the immunohistochemistry of AIMP1/2/3 protein levels with annotated protein-level data, that were downloaded from The Human Protein Atlas (https://www.proteinatlas.org/) for GBM, LGG and normal brain tissue, and reported accordingly. The immunohistochemistry slides have been presented along with annotated protein levels data for gliomas and normal brain tissue in Fig2B&C.

### Epigenetic regulation of AIMP1/2/3 in gliomas

We performed a comprehensive CpG-wide analysis for AIMP1/2/3 genes using TCGA data, where matched transcriptomic data were available. Methylation levels (beta values) for each CpG-site were downloaded for AIMP1/2/3. CpG-site specific methylation levels of AIMP1/2/3 were correlated with their respective gene mRNA expression levels, to identify potential epigenetic regulators of gene expression in TCGA-GBM and TCGA-LGG. Pearson’s correlation coefficients were reported, with p-value<0.05 considered statistically significant. Heatmaps were created with the Seaborn package using Python. Significant correlations were marked with asterisk on the heatmaps, while negative and positive correlations were marked with hues of blue and red, respectively.

### Prognostic effects of AIMP1/2/3 mRNA expression and CpG-level methylation

We analyzed the prognostic effects of AIMP1/2/3 mRNA expression on the overall survival of GBM and LGG patients in TCGA, CGGA, REMBRANDT and GRAVENDEEL datasets. Survival analysis for was conducted using the Kaplan-Meier method, and the “survival” and “survminer” packages in R were utilized to assess the survival effects. Log-rank p-value <0.05 was considered statistically significant. We analyzed the AIMP1/2/3 CpG-level methylation prognostic effects on the overall survival of TCGA-GBM and TCGA-LGG datasets using Cox Proportional Hazards model. Hazard ratios with 95% confidence intervals were reported. Chi-square p-value < 0.05 were considered statistically significant. Bonferroni correction was used for adjusted p-values.

We analyzed the prognostic effects of specific CpG-level methylation of AIMP1/2/3 within subgroups of bevacizumab treated patients only in TCGA-GBM (n=80) and TCGA-Astroctyoma (n=47) using Cox Proportional Hazards model for overall survival and disease-free survival. Both univariate analysis and multivariate analysis adjusting for confounding factors were performed. R-packages “survival” and “forestplot” were used for analysis. Hazard ratios with 95% confidence intervals were reported. Chi-square p-value < 0.05 were considered statistically significant. Bonferroni correction was used for adjusted p-values. Only significant CpGs were visualized in forest plots. For TCGA-Astroctyoma with available treatment response data (n=22), we compared stable versus progressive disease using Student’s t-test (p-value<0.05 considered statistically significant).

### Retrospective analysis of GBM clinical trial datasets: REGOMA and BELOB

We examined the efficacies of anti-angiogenic therapies in recurrent GBM clinical trials, REGOMA and BELOB. The REGOMA trial was a multicenter, open-label, randomized, controlled phase II trial (REGOMA) for investigating the effect of regorafenib in patients with recurrent GBM (n=71 where transcriptomic data were available). The BELOB trial was a phase II, multicenter, randomized clinical trial that evaluated the efficacy of bevacizumab and lomustine in patients with recurrent GBM (n=112 where transcriptomic data were available). We investigated the efficacies of these treatments in AIMP1/2/3 high- and low-mRNA expression subgroups stratified by median expression cut-off values. In each subgroup, we performed the Kaplan-Meier method and Cox Proportional Hazards model analysis using IBM SPSS, with p-value<0.05 considered significant in log-rank and Chi-square p-values, respectively.

### Single-cell transcriptomic and spatial transcriptomic analysis

We previously preprocessed, normalized and concatenated three single-cell transcriptomic cohorts - GSE131928 (n=28), GSE163108 (n=16), and GSE84465 (n=4) [53]. We have also previously annotated the cell-types in these datasets as outlined by Zheng et. al [53]. These annotations were used for cell-type specific comparisons in this work.

Python package ‘scanpy’ was used for spatially visualizing transformed spatial transcriptomic data for AIMP1/2/3. AIMP1/2/3 levels were compared between GBM cell-types (AC-like, OPC-like, MES-like1, MES-like2 and NPC-like) in the spatial transcriptomic cohort and the concatenated single-cell transcriptomic cohort (consisting of three datasets). Violin plots were generated using Python ‘seaborn’ package. Kruskal-Wallis p-value <0.05 was considered statistically significant. Mean and median expression values were calculated for each cell-type to determine cell-types with highest AIMP2 expression. Only AIMP2 showed significant differences across cell-types in both cohorts.

### Statistical Analysis

All statistical analyses were performed using R (version 4.2.2) and Python (version 3.10.9). Data preprocessing and normalization were conducted as per dataset-specific protocols. Student’s t-test was used for pairwise comparisons (e.g., tumor vs. normal tissues, primary vs. recurrent gliomas) with p < 0.05 considered statistically significant. Pearson’s correlation was used to assess associations between AIMP1/2/3 expression and angiogenesis gene signatures, with results visualized via heatmaps using the Seaborn package.

Survival analyses were performed using the Kaplan-Meier method and Cox Proportional Hazards models, utilizing the “survival” and “survminer” R packages. Log-rank tests determined statistical significance (p < 0.05). Cox models were adjusted for relevant clinical covariates, reporting hazard ratios (HR) with 95% confidence intervals (CI). Multiple hypothesis testing was corrected using Bonferroni adjustment where applicable. For the REGOMA and BELOB clinical trials, treatment response was assessed using Kaplan-Meier survival analysis and Cox regression, stratified by median AIMP expression. IBM SPSS Statistics (version 29.0.2.0) was used for these analyses, with log-rank and Chi-square p-values < 0.05 deemed significant.

Single-cell and spatial transcriptomic data were analyzed using the Scanpy Python package. Kruskal-Wallis tests (p < 0.05) compared AIMP expression across GBM cell subtypes, with violin plots generated via Seaborn.

## Ethics Statement

Details of all available data are provided in the manuscript. No institutional review board or ethics committee approval was required for this manuscript.

## Author contributions

Conceptualization, Investigation, Visualization and Writing—original draft preparation: H.N

Methodology and Data curation: H.N and Y.Z

Supervision: O.G

Writing—review and editing: H.I and O.G

## Data Availability

All data used in this study are publicly available from previously published sources. No new datasets were generated or analyzed beyond those obtained from public repositories. Details on data sources and accession numbers are provided in the Methods section.

## Funding Statement

Research reported here was further supported by the National Cancer Institute (NCI) under awards: R01 CA260271. The content is solely the responsibility of the authors and does not necessarily represent the official views of the National Institutes of Health.

## Notes

### Competing Interest Statement

The authors have declared no competing interest.

