## Supplementary File for "Response to anti-angiogenic therapy is affected by AIMP protein family activity in glioblastoma and lower-grade gliomas"

**SUPPLEMENTARY FIGURES**

**Supplementary Figure 1**

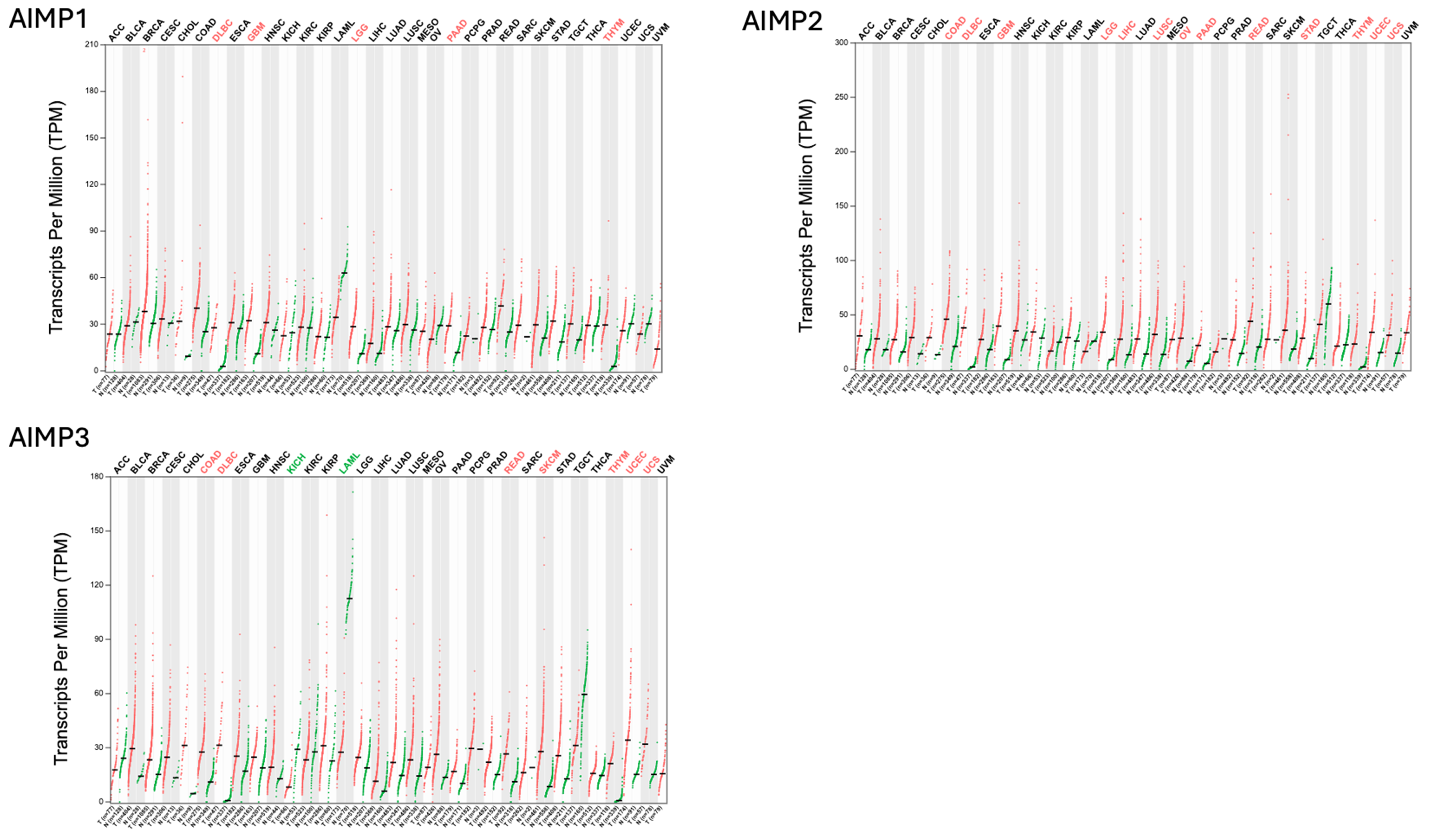

**Supplementary Figure 1.** Pan-cancer comparison of AIMP1/2/3 mRNA expression levels between tumor (TCGA) versus normal tissue (GTEx). Red label indicates significant association with higher expression in the tumor; Green label indicates significant association with higher expression in the normal tissue (p<0.05).

**Supplementary Figure 2**

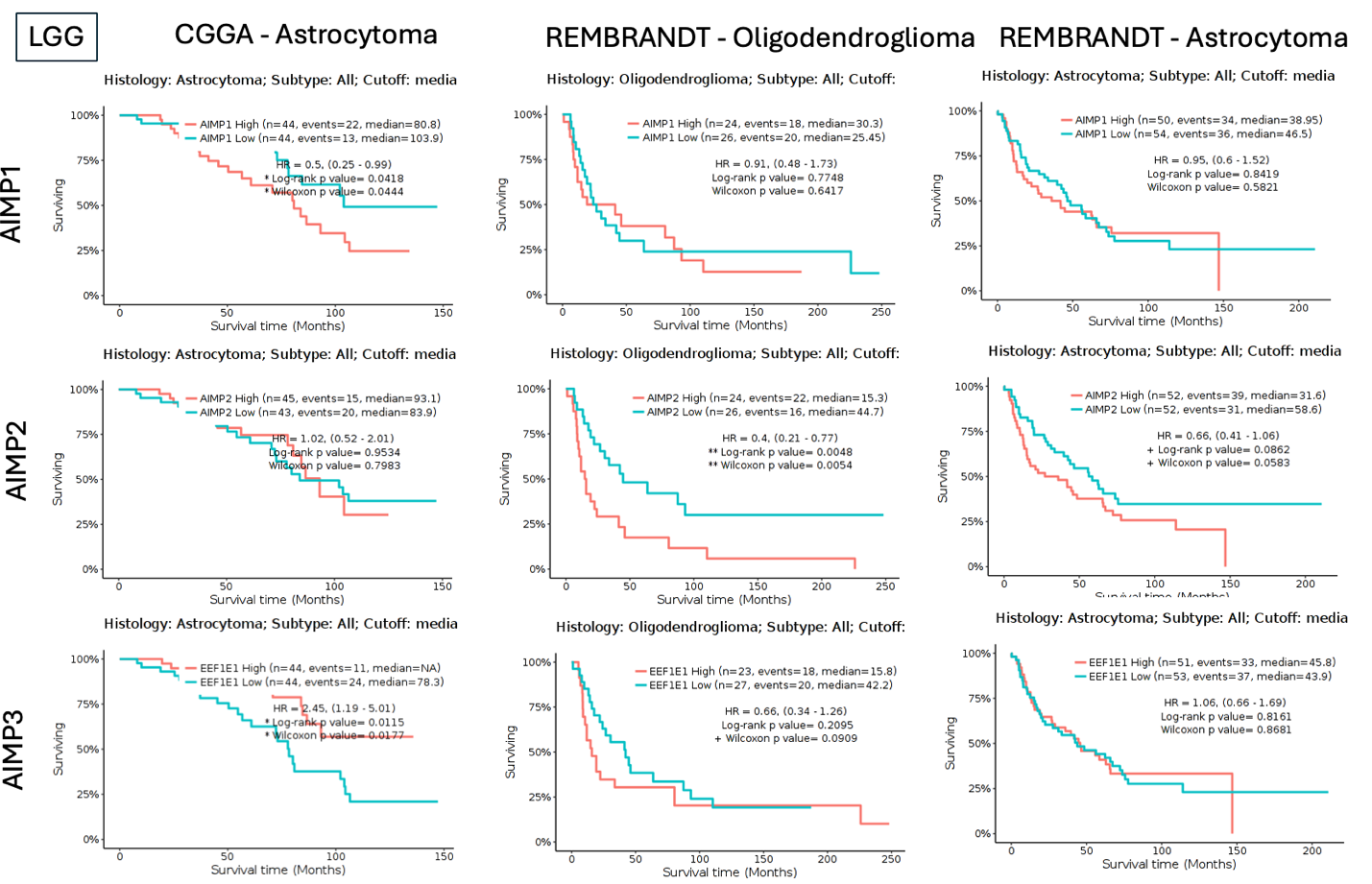

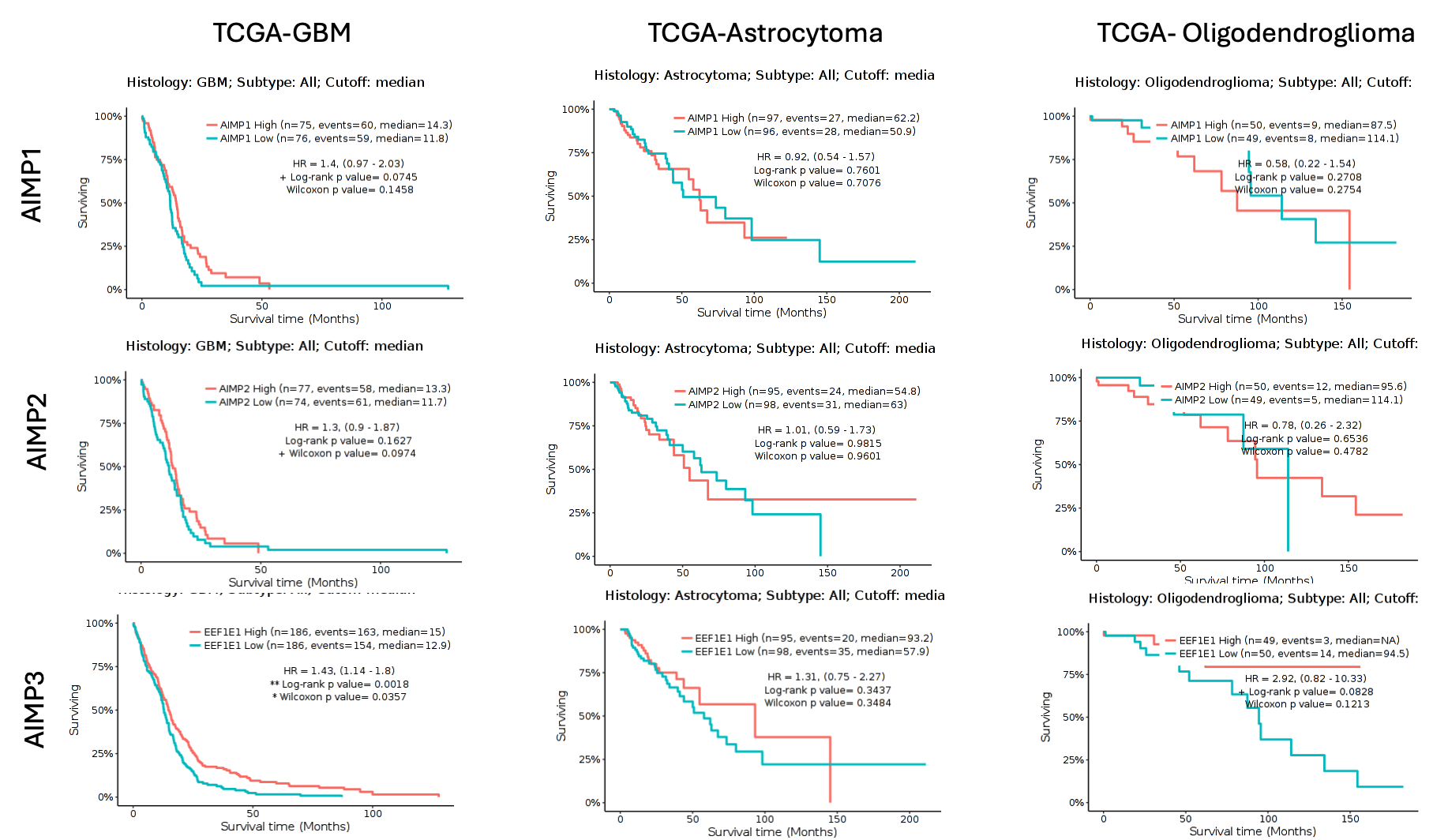

**Supplementary Figure 2.** Kaplan-Meier Survival Curves depicting prognostic effects of AIMP1/2/3 mRNA expressions in CGGA, TCGA, and REMBRANDT cohorts of gliomas. Red=high-expression green=low-expression based on median-cutoff; log-rank p-value<0.05 is considered significant.

**Supplementary Figure 3**

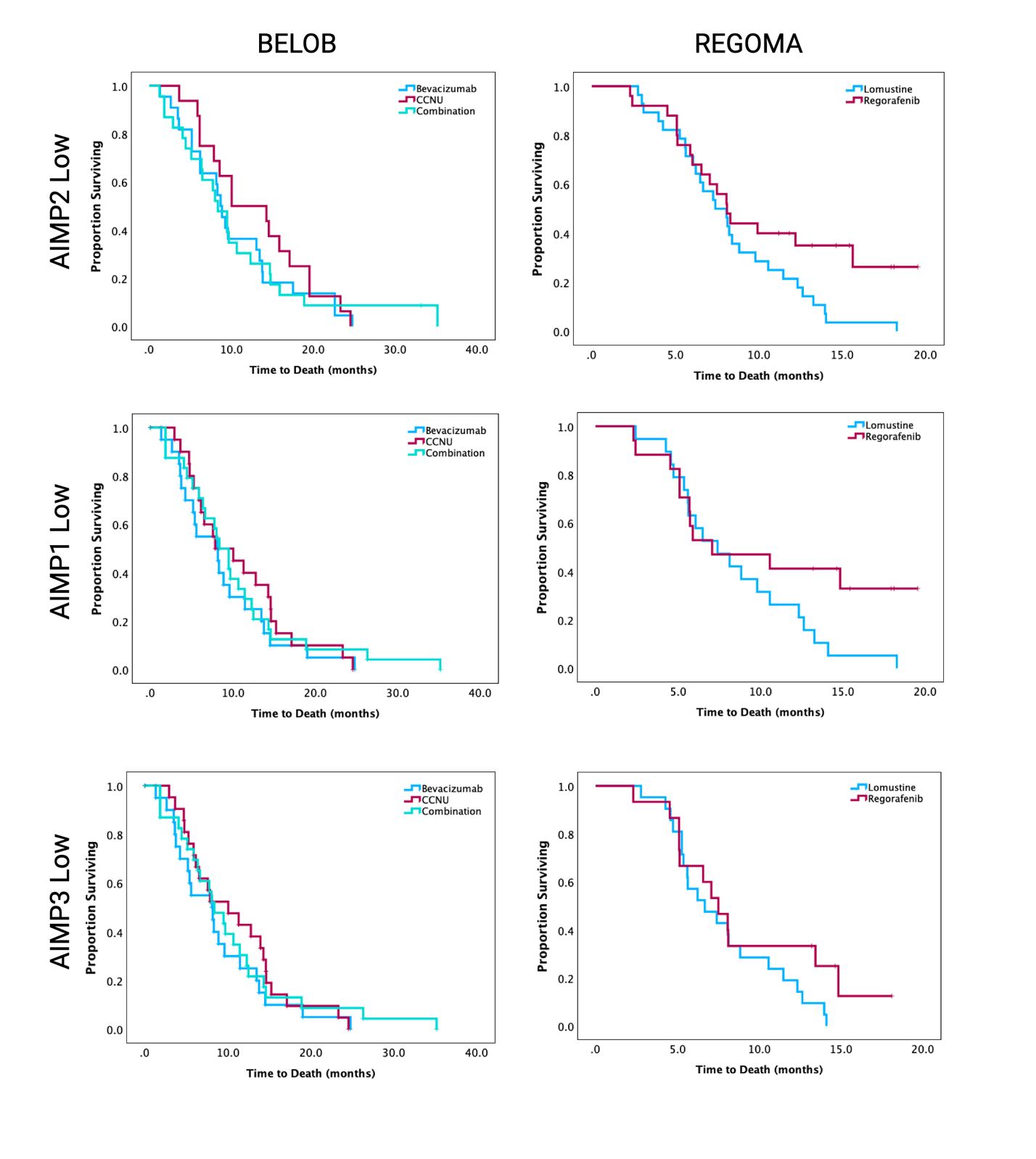

**Supplementary Figure 3. Low AIMP mRNA expression subgroups are not responsive to anti-angiogenic therapies.** Kaplan-Meier survival analysis on retrospective clinical trials of recurrent GBM (REGOMA and BELOB trials). Low expression sub-groups are stratified by median mRNA expressions of AIMP1/2/3. Log-rank p-value<0.05 is considered significant.

**Supplementary Figure 4**

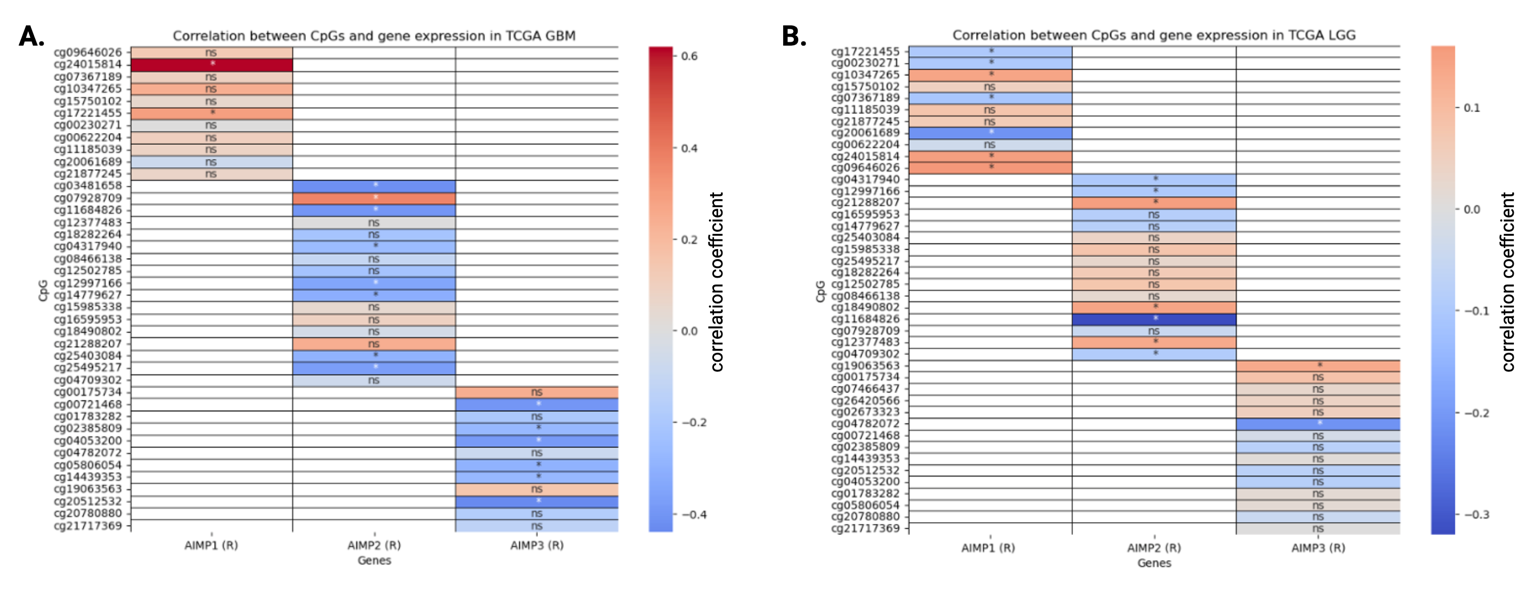

**Supplementary Figure 4.** Correlation between specific AIMP1/2/3 CpG site methylation levels and their respective mRNA expression levels in **(A)** TCGA-GBM and **(B)** TCGA-LGG. *asterisk represents significant correlation (p-value<0.05). The colormap indicates blue hues=negative correlation and red hues=positive correlation.

**Supplementary Table 1 : Prognostic AIMP1/2/3 CpG-sites in TCGA GBM and TCGA Astrocytoma**

|  |  | **GBM OS** |  |  |  |
| --- | --- | --- | --- | --- | --- |
| **AIMP1** | coef | Hazard Ratio | se(coef) | z | p-value |
| cg20061689 | -28.19 | 5.72E-13 | 13.69 | -2.05 | 0.039 |
| cg07367189 | -18.14 | 1.32E-08 | 8.145 | -2.22 | 0.0258 |
| **AIMP2** |  |  |  |  |  |
| cg11684826 | 2.33 | 10.35 | 1.137 | 2.055 | 0.0398 |
|  |  | **GBM DFS** |  |  |  |
| **AIMP2** |  |  |  |  |  |
| cg11684826 | 3.36 | 29.00 | 1.35 | 2.49 | 0.0127 |
|  |  | **Astrocytoma DFS** |  |  |  |
| **AIMP3** |  |  |  |  |  |
| cg05806054 | 3.15 | 23.54 | 1.17 | 2.69 | 0.0069 |
| cg01783282 | 2.50 | 12.18 | 1.21 | 2.06 | 0.0393 |

**Supplementary Figure 5**

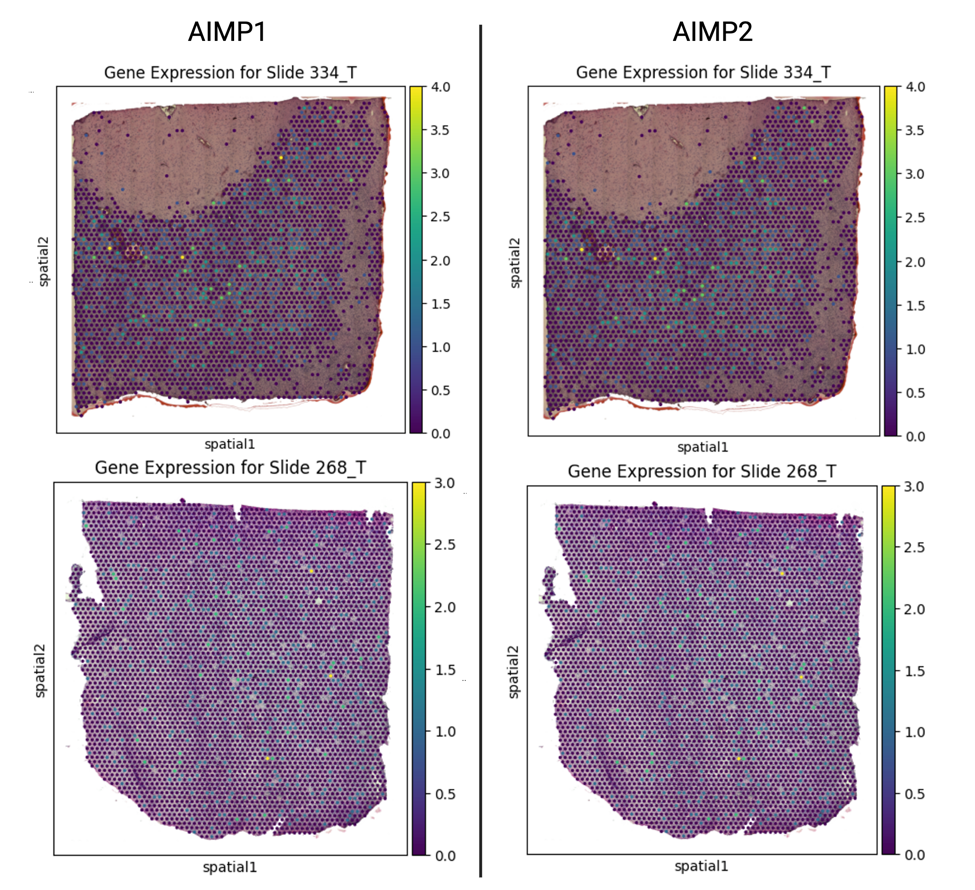

**Supplementary Figure 5. Spatial expression of AIMP1 and AIMP3 in glioblastoma (GBM) tumors.** Spatial distribution of AIMP1 and AIMP3 expression across tumor tissues in two representative GBM slides (i: Slide 334_T, ii: Slide 268_T) analyzed using a spatial transcriptomic dataset. The heatmaps display homogeneous expressions of AIMP1 and AIMP3 with varying intensity from low (purple) to high (yellow), indicating the absence of localized expression hotspots.
